# Alternative splicing and environmental adaptation in house mice

**DOI:** 10.1101/2023.06.23.546335

**Authors:** David N. Manahan, Michael W. Nachman

**Affiliations:** Department of Integrative Biology and Museum of Vertebrate Zoology University of California, Berkeley Berkeley, CA 94720, USA

## Abstract

A major goal of evolutionary genetics is to understand the genetic and molecular mechanisms underlying adaptation. Previous work has established that changes in gene regulation may contribute to adaptive evolution, but most studies have focused on mRNA abundance and only a few studies have investigated the role of post-transcriptional processing. Here, we use a combination of exome sequences and short-read RNA-Seq data from wild house mice (*Mus musculus domesticus*) collected along a latitudinal transect in eastern North America to identify candidate genes for local adaptation through alternative splicing. First, we identified alternatively spliced transcripts that differ in frequency between mice from the northern-most and southern-most populations in this transect. We then identified the subset of these transcripts that exhibit clinal patterns of variation among all populations in the transect. Finally, we conducted association studies to identify *cis*-acting splicing quantitative trait loci (*cis*-sQTL), and we identified *cis*-sQTL that overlapped with previously ascertained targets of selection from genome scans. Together, these analyses identified a small set of alternatively spliced transcripts that may underlie environmental adaptation in house mice. Many of these genes have known phenotypes associated with body size, a trait that varies clinally in these populations. We observed no overlap between these genes and genes previously identified by changes in transcript level, indicating that alternative splicing and changes in mRNA abundance may provide separate molecular mechanisms of adaptation.

## Introduction

Nearly 50 years ago, King and Wilson (1975) argued that the proteins of humans and chimpanzees were so similar that changes in gene regulation might underlie much of organismal evolution. More recent work has provided strong evidence in support of the idea that differences in gene expression underlie adaptation (e.g. Wray et al. 2003; Jones et al. 2012; Fraser 2013; Mack et al. 2018). Most of this work has focused on differences in mRNA abundance, but gene regulation involves many steps, including transcription, post-transcriptional modifications, translation, and post-translational modifications. Due to challenges in measuring post-transcriptional modifications, these have been less studied as mechanisms for evolution and local adaptation (but see Artieri & Fraser 2014; Yablonovitch et al. 2017; Steward et al. 2022). One important post-transcriptional modification is alternative splicing. During mRNA splicing, introns are typically removed and some or all of the exons in a gene are joined together in a processed mRNA, or isoform. While some exons may be constitutively spliced in, other exons may be alternately spliced in or out to generate numerous unique mRNA isoforms from a single gene (Fig. 1A). Through alternative splicing of exons and introns, protein domains can also be included or excluded in the translated protein. Mutations in regions for *cis*- and *trans*-regulatory control of splicing can therefore alter protein structure and function and generate different phenotypes (Manning et al. 2017). These phenotypes, in turn, present opportunities for selection to drive divergence between populations.

**Fig 1.**
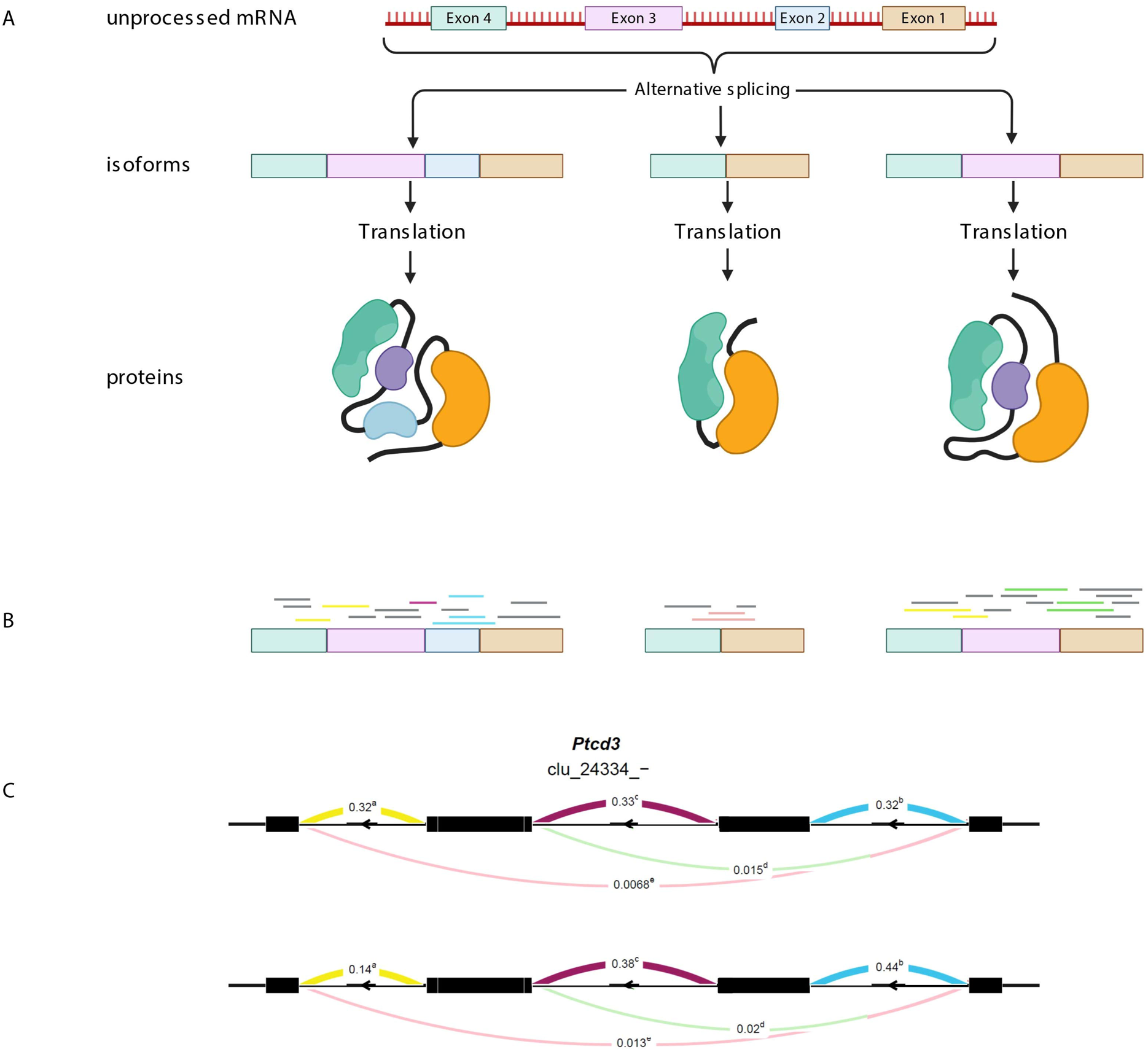
Alternative splicing generates a diversity of protein folds and functions from a single gene. **A** An overview of alternative splicing. Following transcription of a gene, exons are joined together in different combinations to produce different isoforms. These isoforms are related, but by including and excluding exons in the processed mRNA, protein folding can be changed and entire domains can be included or excluded in the translated product. Created with Biorender.com. **B** Mapping RNA-Seq short reads to mRNA isoforms. Short-read expression data is unable to reliably capture isoform diversity. Instead of defining alternative splicing by quantifying isoforms, short-reads can be used to define alternative splicing events. Reads that map to two different loci (split reads) capture splicing events between two junctions. **C** Leafcutter (Li et al. 2018) uses split reads counts to quantify the frequency of splicing between two exons, as seen in this cluster diagram. For example, the blue split reads indicate that exon 1 and 2 and spliced together at a frequency of 0.32 and 0.44 in two different populations. The change in percent splice in (ΔPSI) is 0.12.

Alternative splicing has been shown to underlie adaptation over short timescales in diverse animals from vertebrates to insects (e.g. Jarosch et al. 2011; Ceinos et al. 2018; reviewed in Wright et al. 2022). For example, alternative splicing underlies differential heat sensitivity in vampire bats. Both vampire bats (*Desmodus rotundus)* and closely related fruit bats (*Carollia brevicauda*) express full length *TRPV1*, a heat sensitive ion channel. Full length *TRPV1* is activated at temperatures higher than 38 °C to detect noxious heat. *TRPV1* isoforms expressed in the specialized pit organs of *D. rotundus* are alternatively spliced to include an exon containing a premature stop codon that is translated into a truncated protein (Gacheva et al. 2011). Truncated *TRPV1* is activated at 30 °C and improves sensitivity for detecting homeothermic prey for blood feeding. Alternative splicing has also been shown to facilitate adaptation by producing isoforms with loss-of-function phenotypes. Threespine sticklebacks (*Gasterosteus aculeatus*) are well known to have marine and freshwater ecotypes, with a reduction of spines in freshwater ecotypes. Marine sticklebacks produce full length isoforms of the homeodomain transcription factor *MSX2A* while freshwater sticklebacks produce a high proportion of non-functional *MSX2A* by splicing out an exon encoding a DNA binding domain (Howes et al. 2017). Transgenic expression of marine isoforms in freshwater sticklebacks increased spine length, suggesting that alternative splicing can facilitate adaptation by producing non-functional proteins.

The Western European house mouse (*Mus musculus domesticus*) provides a useful system for exploring the potential role of alternative splicing in local adaptation. Native to the Mediterranean region and Western Europe, house mice were spread around the world in association with humans in the last few hundred years (Morgan et al. 2022). In this short time they have adapted to a wide range of different environments through changes in morphology, physiology, and behavior. For example, mice in Eastern North America show clinal patterns of variation in body size, with larger mice in colder environments consistent with Bergmann’s Rule (Bergmann 1847), and these differences persist in a common lab environment (Lynch 1992). In eastern North America, mice from colder environments also build bigger nests, differ in metabolic traits, and show population-level differences in gene expression compared to mice from warmer environments when reared in a common laboratory environment (Phifer-Rixey et al. 2018). Moreover, patterns of genomic variation in mice sampled along a latitudinal transect from Florida to New York identified specific candidate genes underlying adaptive phenotypic differences (Phifer-Rixey et al. 2018). Most of these candidate genes did not harbor non-synonymous variants, suggesting that much of the response to selection is driven by changes in gene regulation. By identifying overlap between loci exhibiting clinally varying patterns of gene expression, genes with *cis*-eQTL, and loci showing signatures of selection, Mack et al. (2018) were able to identify a small number of genes responsible for adaptive differences in body size.

In this study we expand on previous work in house mice by exploring the role of splice variants in environmental adaptation. In particular, we analyzed RNA-Seq data from liver in 50 wild-caught mice sampled in five populations along a latitudinal transect (Fig. 2). First, we compared mice from the ends of this transect to identify genes showing population-level differences in alternative splicing. Next, we asked which of these genes showed clinal patterns of alternative splicing as might be expected if selection is driving differences. Third, we looked for associations between SNPs near each gene and splice variants to identify *cis*-acting splicing quantitative trait loci (*cis*-sQTL). Fourth, we explored the overlap among these loci and those showing signatures of selection from previous work (Phifer-Rixey et al. 2018) to identify a small set of candidate splice variants underlying environmental adaptation. Finally, we compared these results on alternative splicing with similar analyses conducted on patterns of mRNA level (Mack et al. 2018). We discovered five genes that showed differential splicing between mice sampled from the ends of this transect, clinal patterns of splicing variation across all five populations, and *cis*-sQTL that overlapped with previously identified targets of selection (Phifer-Rixey et al. 2018). We observed little to no overlap between genes showing alternative splicing and those showing overall differences in mRNA level.

**Fig 2.**
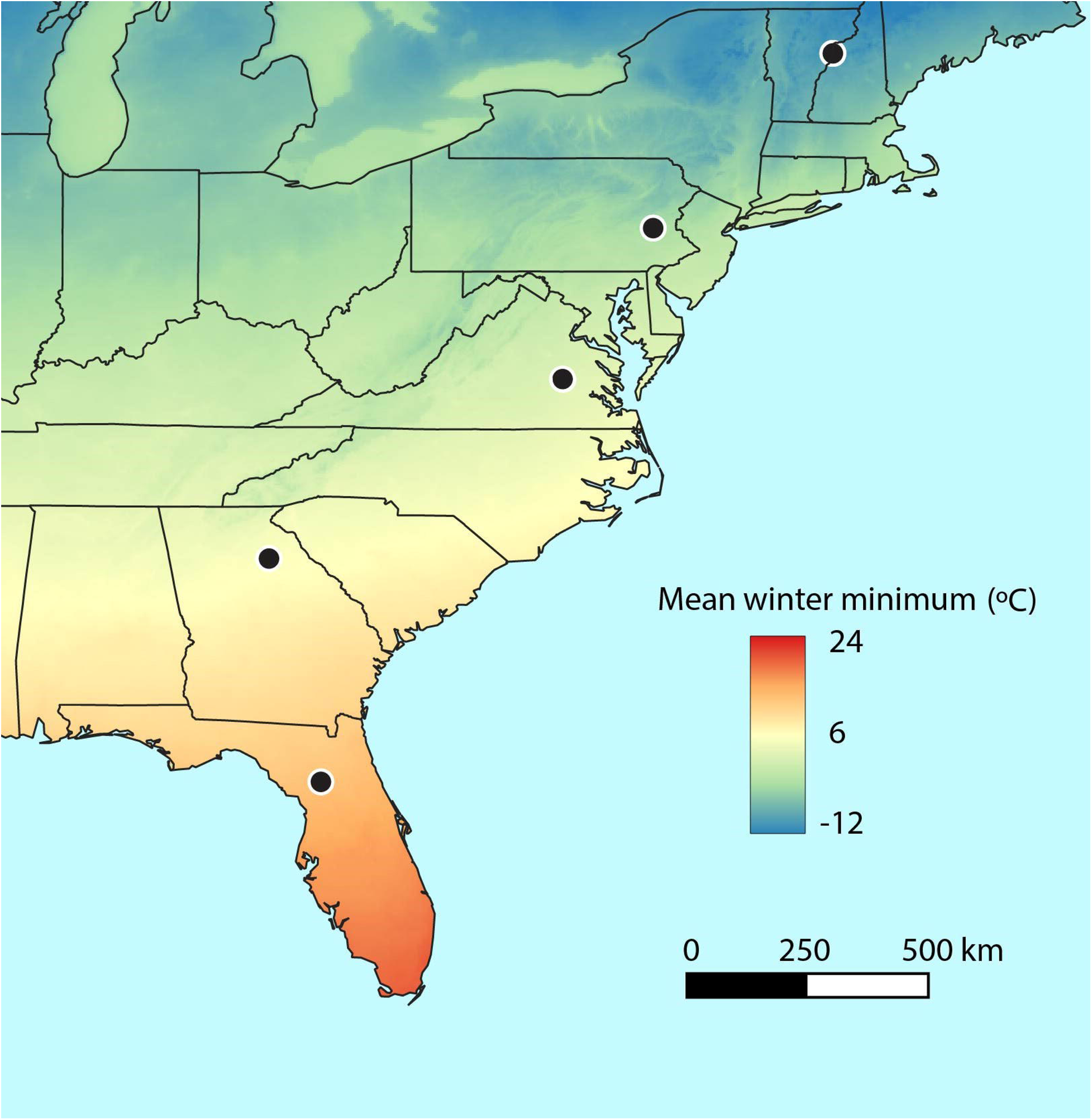
Sampling distribution of house mice along a latitudinal cline. Five populations were included for study: New Hampshire/Vermont, Pennsylvania, Virginia, Georgia, and Florida. Latitude was used as a proxy for bioclimatic variables, including mean winter minimum temperature (climate data is from Bioclim.org).

## Materials and Methods

### Sampling and mRNA-sequencing

RNA-Seq data were from Mack et al. (2018) (NCBI Bioproject: PRJNA407812). *Mus musculus domesticus* were collected from five populations along a latitudinal transect in the eastern United States (New Hampshire/Vermont, Pennsylvania, Virginia, Georgia, and Florida; Fig. 2). Mice were sacrificed in the field, and liver tissues were extracted and stored in RNAlater at 4°C overnight before being frozen at -80°C. RNA was extracted with Qiagen’s RNeasy Mini kit.

RNA-Seq libraries of 100-base pair paired-end reads were generated for each population on the Illumina HiSeq 4000 platform.

### mRNA-sequence data processing

We trimmed mRNA-sequence reads from Mack et al. (2018) with TrimGalore! version 0.5.0 and we mapped these reads with STAR version 2.7.9a (Spliced Transcripts Alignment to a Reference) to the GRCm39 reference genome (Dobin et al. 2013). STAR has improved ability to identify annotated exon junctions and contains the `twopassMode` option that uses exon junctions found in the first run of mapping as annotations in a second run of mapping. This option helps to identify unannotated exon junctions. The output from STAR consists of mapped reads in .bam files which we then converted into .junc files containing the mapped reads that span exon junctions using Regtools junction version 0.5.2 (Cotto et al. 2020).

### Exon clustering for alternative splicing and differential splicing analysis

We used Leafcutter to identify clusters of alternatively spliced exons from .junc files (Li et al. 2018). Junctions were identified from the subset of RNA-Seq data that are split reads (i.e., reads that map to two unique locations in a gene). All junctions that shared either a splicing donor or acceptor site are considered part of the same cluster. These clusters represent the set of exons that can be alternatively spliced with one another. Genes typically had multiple alternatively spliced clusters (Fig. 3A). We also used Leafcutter to detect differentially spliced exon clusters between the northernmost (New Hampshire/Vermont) and southernmost (Florida) populations from the latitudinal transect using a Dirichlet-multinomial generalized linear model. Testing for differential splicing between clusters has improved sensitivity over tests for differential splicing between exons tested individually. Differential splicing was measured as the ΔPSI (change in percent spliced-in) between the two populations for each exon in a cluster (Fig. 3B). Leafcutter also calculates principal components explaining variation in junction counts between populations.

**Fig 3.**
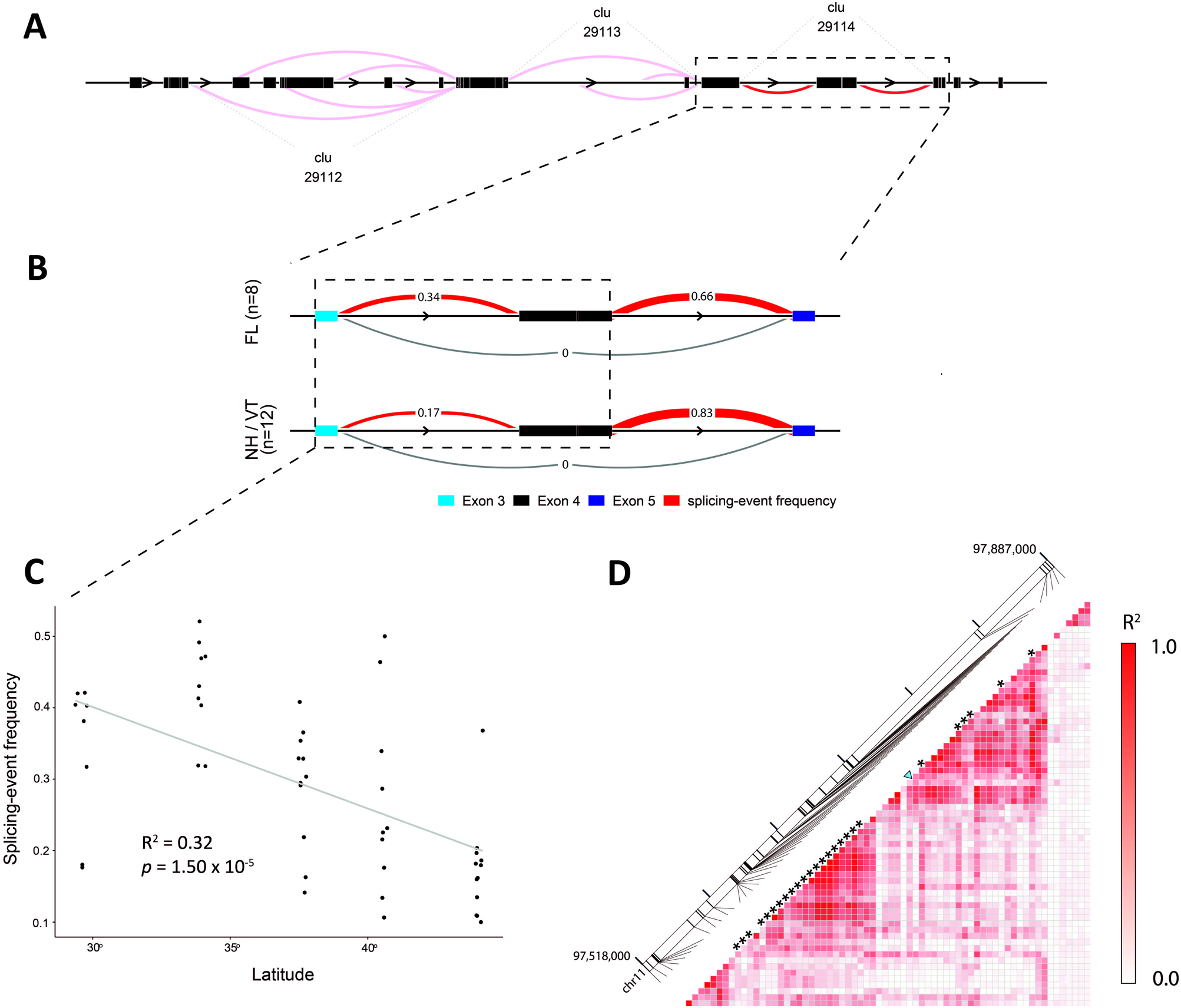
Identification of candidates for adaptive splicing. *Lasp1* is a candidate for adaptive splicing among house mice in eastern North America. **A** Overlapping splicing donor and acceptor sites were used to identify clusters of alternatively spliced exons. The *Lasp1* gene contains three differentially spliced clusters (arbitrarily named 29112, 29113, and 29114). **B** Differential splicing was assessed between the northernmost (New Hampshire/Vermont) and southernmost (Florida) populations on the latitudinal cline. *Lasp1* had one cluster (29114) that exhibited differential splicing (FDR < 0.05). Cluster 29114 contained exons three, four, and five of *Lasp1*. Leafcutter measures differentially splicing in terms of the change in percent-spliced in (ΔPSI) between populations. In *Lasp1*, exon 3 and 4 are spliced in at an average frequency of 0.34 and 0.17 between Florida and New Hampshire/Vermont populations, respectively and the ΔPSI for those exons is 0.17 **C** For each differentially spliced cluster between New Hampshire/Vermont and Florida, the splice-in frequency for the pair of exons with the highest ΔPSI was fitted to a linear model with latitude. The frequency of splicing between exon 3 and 4 of *Lasp1* is correlated with latitude (*p* = 1.50 x 10^-5^, R^2^ = 0.32). **D** *cis*-sQTL for each clinally varying splicing event were identified with a linear-mixed model. Linkage disequilibrium between *cis*-sQTL (blue triangle) and LFMM outliers (asterisks) was calculated based on correlation.

### Identification of clinally varying splicing events

For each differentially spliced cluster between New Hampshire/Vermont and Florida, the exon with the great ΔPSI was selected to test for clinal variation since it is likely that the splice-in-frequency of exons in the same cluster are not independent of each other. We used linear regression models of splice-in-frequencies on latitudes from all five populations to identify differentially spliced exons whose frequency varies clinally across latitude (*p* < 0.05, ΔPSI > 0.10). Linear regression models were calculated in R (version 4.2.3).

### Mapping and variant calling on exome-capture data

We mapped reads from the exome capture data of Phifer-Rixey et al. (2018) to GRCm39 with Bowtie2 v2.4.1 (Langmead & Salzberg 2012). Variant calling was performed with Genome Analysis Toolkit v4.1 (GATK) HaplotypeCaller (McKenna et al. 2010). The Base Recalibrator tool in GATK was used to identify and remove sequencing errors. Alleles missing from 30% of individuals and with a minor allele frequency less than 10% were removed from analysis. The filtered set of SNPs was then used in association studies to detect *cis*-sQTL.

### Cis-sQTL discovery and overlap with signals of selection

We used a generalized linear mixed model implemented in the program GEMMA version 0.98.3 (Genome-wide Efficient Mixed Model Association) to identify *cis*-sQTL (Zhou et al. 2012). This model calculates a relatedness matrix among genotypes and includes the relatedness matrix as a covariate when testing for association between SNPs and splice variants. Associations were tested for each gene that contained a differentially spliced cluster with an exon whose splice-in frequency varied clinally. SNPs found within 200 kb of each gene were tested for associations with splice variants. We applied the Benjamini-Hochberg correction to each *p*-value, and SNPs with adjusted *p*-values <0.05 were considered significant.

To identify splice variants under selection, we looked for overlap between *cis*-sQTL and SNPs with signatures of selection from Phifer-Rixey et al. (2018). In that study, Latent-Factor Mixed Modeling (LFMM) was used to detect signatures of selection because the method accounts for covariance between environmental and genetic variation (Frichot et al. 2013). Colocalized *cis*-sQTL and LFMM outliers were identified, and for genes without colocalized SNPs, linkage disequilibrium was calculated between *cis*-sQTL and LFMM outliers using PLINK and Haploview (Purcell et al. 2007; Barrett et al. 2005).

## Results

### Differential splicing in wild-house mice

After mapping with STAR, we identified ∼1.8 billion RNA-Seq short-reads across all individuals, and ∼720 million (39%) of those were split reads, averaging ∼14 million split reads per sample. Clusters were only considered for differential splicing if each junction was supported by at least 10 reads. Principal component analysis of all five populations using junction count data showed that individuals did not cluster by population, although individuals clustered more tightly in populations from warmer climates (Supplementary Fig. 1). The first two principal components explained 3.6% and 2.9% of the variance respectively. The first 10 principal components accounted for 27.1% of the variation and none resulted in population-level clustering.

We used Dirichlet-multinomial generalized linear model implemented in Leafcutter to test for differential splicing between the New Hampshire/Vermont and Florida populations at the cluster level. Differential splicing was measured as the difference between two populations in the proportion of spliced-in exons (Fig. 3B). This test identified 168 differentially spliced exon clusters in 152 genes between the two populations (Supplementary Table 1; *p* <0.05, Benjamini-Hochberg correction).

### Clinal patterns of alternative splicing

Patterns of genetic variation among the five sampled populations do not show isolation-by-distance (Phifer-Rixey et al. 2018). Because of this, traits that co-vary with environmental gradients may be indicative of selection. We tested for clinal patterns of variation among differentially spliced clusters using latitude as a variable because latitude is strongly correlated with many climatic variables. For example, latitude shows a strong negative correlation with the first principal component summarizing climatic variables from the WorldClim database (Hijmans et al. 2005) (Pearson’s r = -0.99, *p* < 0.0006, Phifer-Rixey et al. 2018). We considered clusters with a maximum |ΔPSI| less than 0.10 to have a very small effect, and these were excluded from the analysis. Of the 152 genes with differentially spliced exons, we identified 44 exons in 41 genes that were correlated with latitude and showed |ΔPSI| > 0.10 (Fig. 3C; Supplementary Table 2, *p* < 0.05). A gene ontology overrepresentation test showed no significant enrichment for either molecular function, biological process, or cellular component in this set of 41 genes (Panther overrepresentation test, FDR < 0.05).

### Identification of cis-sQTL and overlap with signals of selection

To identify *cis*-sQTL for each of the 44 clinally varying exons, we used a linear mixed model to test for associations between all variants within 200 kb of the acceptor site and the splice-in frequency of the corresponding exon, with relatedness between individuals included as a covariate. In total, 829 variants from the exome-capture data of Phifer-Rixey et al. (2018) were tested. Out of the 44 clinally varying exons, 24 exons in 21 genes had at least one *cis*-sQTL (Supplementary Fig. 3, -log_10_(*p*) > 1.30, *p* < 0.05, Benjamini-Hochberg correction).

Next, we looked for overlap between the 22 *cis*-sQTL and signals of selection based on LFMM outliers with |Z-scores| > 2 across all genes from Phifer-Rixey et al. (2018). Five of the 22 *cis*-sQTL overlapped with these LFMM hits at the gene level, and this amount of overlap is no more than expected by chance (Supplementary Fig. 3, permutation test, *p* = 0.601). To assess whether these five genes contained *cis-*sQTL and LFMM outliers on the same haplotype, we calculated linkage disequilibrium among all SNPs in the 200 kb window for each gene (Fig. 3D, Supplementary Fig. 4). Each candidate either had colocalized *cis*-sQTL and LFMM outliers or had *cis*-sQTL and LFMM outliers in the same haploblock. These five genes with *cis*-regulatory variants underlying clinal patterns of splicing are strong candidates for adaptive evolution mediated by alternative splicing. Notably, most of these five genes are associated with phenotypes known to distinguish mice from different latitudes (Phifer-Rixey et al. 2018). For example, three of the five genes are associated with differences in growth (Table 1) and mice from different latitudes are known to differ in body size and growth rates.

**Table 1.**
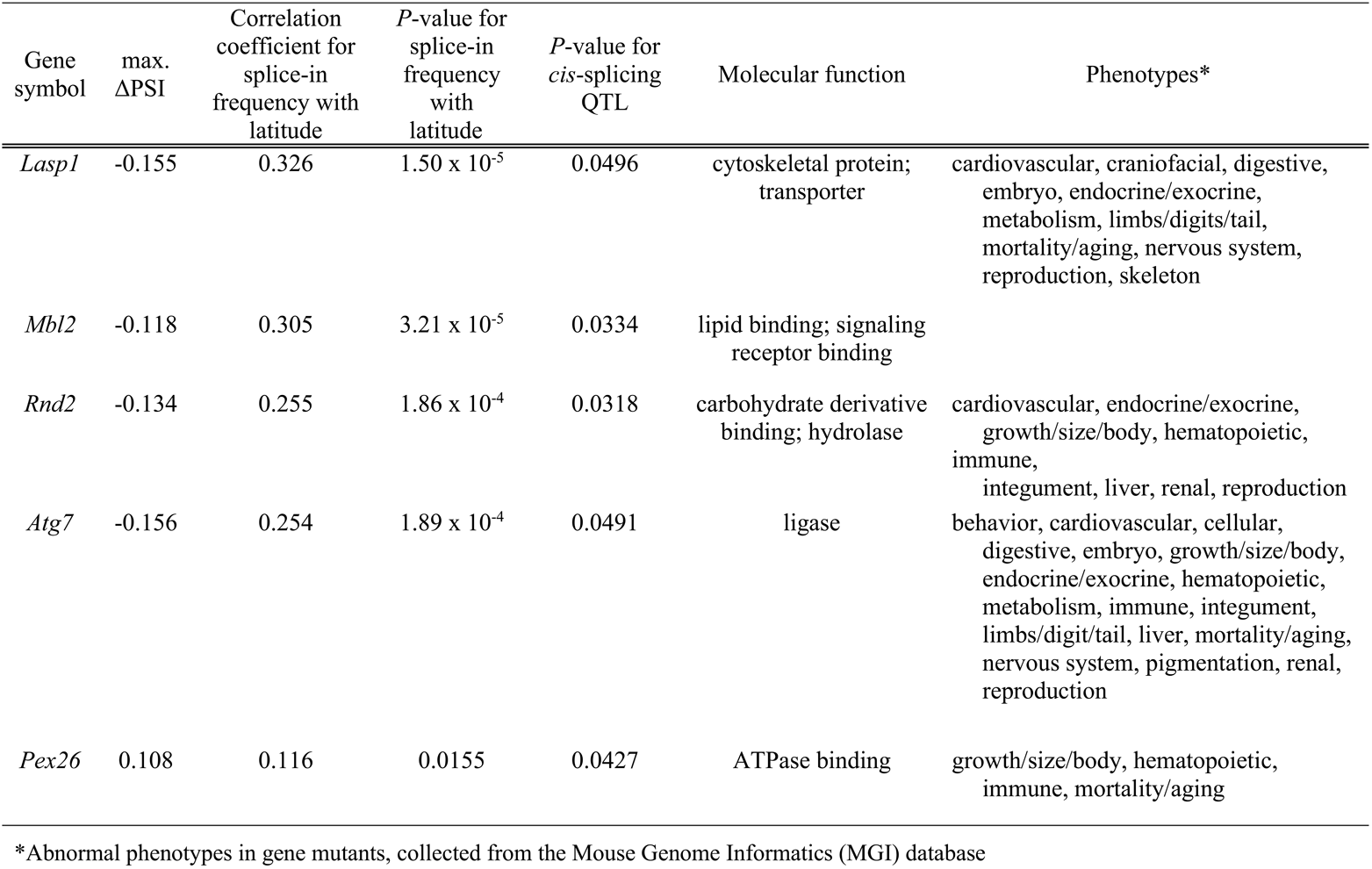
Candidates for adaptive splicing in house mice in eastern North America.

### Comparison between alternative splicing and mRNA abundance

We compared patterns of alternative splicing to the patterns of mRNA abundance (i.e. gene expression) from Mack et al. (2018) (Table 2). In comparisons between the two populations at the ends of this transect, 152 genes showed differential splicing and 458 genes showed differential transcription, with only 6 genes in common. Of these, 41 genes showed clinal patterns of alternative splicing and 274 genes showed clinal patterns of mRNA abundance, with no overlap between these sets of genes. Thus, there is little overlap between genes that show alternative splicing and those that show differences in mRNA abundance. Of the genes with clinal patterns of mRNA abundance, five of those with a *cis*-eQTL overlapped with a LFMM outlier, while for genes showing clinal patterns of splicing variation, five of those with a *cis*-sQTL overlapped with a LFMM outlier, suggesting that a similar proportion of *cis*-eQTL and *cis*-sQTL are targets of selection (Table 2).

**Table 2.**
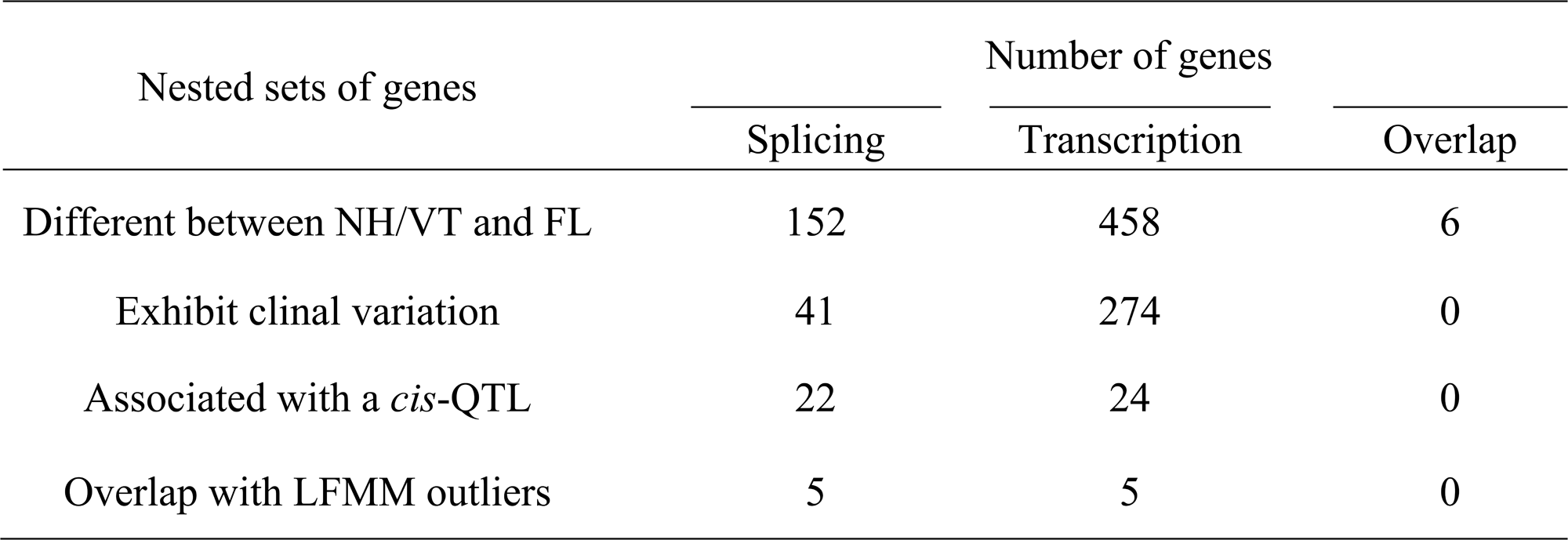
Comparison between splicing and transcript abundance.

## Discussion

We used short-read RNA-Seq data from five populations of house mice sampled along a latitudinal transect to study the role of alternative spicing in environmental adaptation. We found genes that differed in the frequency of alternatively spliced exons in comparisons between mice sampled from the ends of this transect. A subset of those genes showed clinal patterns of splicing variation, and a further subset harbored *cis*-sQTL that overlapped with previously identified targets of selection. Notably, these genes are associated with phenotypes that distinguish mice from the ends of the transect.

The impact of post-transcriptional processing on adaptation is a small but growing field of research. Many previous studies have focused on a particular phenotype associated with alternative splicing in a single gene (Gacheva et al. 2011; Jarosch et al 2011; Howes et al. 2017; Ceinos et al. 2018). In contrast, we looked for signatures of adaptive, alternative splicing genome-wide. Phifer-Rixey et al. (2018) assayed the phenotypes of house mice collected from the same populations at the ends of the latitudinal transect (i.e. New Hampshire/Vermont and Florida). In addition to differences in nest-building behavior, activity, and blood chemistry, mice from these populations differ in body size. The larger body size of mice in New Hampshire/Vermont compared to mice from Florida reflects adaptation to colder environments and is consistent with Bergmann’s Rule. Three of the five candidate genes identified in this study (*Rnd2*, *Atg7*, and *Pex26*) are known to affect growth and body size, suggesting potential links between alternative splicing and adaptive phenotypic change. In addition to these five genes, the other genes showing clinal variation in splicing (but not harboring a *cis-*sQTL) might also underlie adaptive differences. Such genes may reflect downstream effects of *trans*-acting mutations elsewhere in the genome. Ten of the 41 genes with clinal variation in splicing are also known to be associated with growth and body size in house mice.

Heritable variation in gene expression is a well-known source of adaptive evolution but has been primarily studied in the context of transcript abundance. Nonetheless, there is an increasing recognition of the importance of post-transcriptional regulation, and a handful of studies have used genome-wide short-read RNA-Seq to study mRNA abundance in conjunction with alternative splicing in the context of local adaptation (Huang et al. 2021; Jacobs & Elmers 2021; Carruthers et al. 2022; Singh et al. 2022; Steward et al. 2022; Huang et al. 2023). Several patterns have emerged from these studies. First, alternative splicing seems to play an important role in local adaptation in distantly related species, such as insects and vertebrates. Second, *cis*-sQTL have been identified that underlie splicing differences between locally adapted populations. Third, in most cases, there seem to be fewer differentially spliced genes than differentially transcribed genes (but see Carruthers et al. 2022), although this may be an artefact arising from differences in methodology as discussed below. Fourth, most studies find that the genes involved in alternative splicing and alternative transcription are not the same (but see Singh et al. 2022). Fifth, most studies have used pairwise comparisons between diverged populations but have not studied splicing variants along an environmental gradient, leaving open the possibility that some of the observed differences are due to stochastic processes. Our results support and build upon the findings from these previous studies. We observed many genes showing differential splicing between populations in different environments and a substantial fraction also had a *cis*-sQTL. We also found little overlap between genes showing differences in transcription and those showing differences in splicing. Unlike previous studies, we explored variation along an environmental gradient, identifying 41 genes where splicing variants were correlated with latitude. Variation along environmental gradients is consistent with local adaptation as a response to spatially varying selection (Endler 1977). Moreover, by identifying *cis*-sQTL that overlap with targets of selection, we identified a small subset of genes that are strong candidates for adaptive alternative splicing.

By using short-read RNA-Seq data, we were able to directly compare alternative splicing and mRNA abundance. Using these same data, Mack et al. (2018) identified 458 differentially transcribed genes between New Hampshire/Vermont Florida while we identified 152 differentially spliced genes between these populations (Table 2). Differences in the methods and power of these two studies make it difficult to compare the total number of genes identified using each approach. For example, short-read data can only identify alternative splicing via split reads and these comprise only 41% of the total reads from the RNA-Seq data. However, the *proportions* of genes in subsequent analyses (i.e. clinally varying, harboring a *cis*-QTL, and overlapping with LFMM outliers) can be directly compared between these two studies (Table 2). Only six genes showed both differential transcription and differentially splicing. Furthermore, there was no overlap in the genes identified for transcription and splicing in any of the other analyses. These results are consistent with previous work comparing adaptation through transcription and splicing. Jacobs and Elmer (2021) also found low overlap between these two mechanisms in the context of adaptation among benthic and pelagic ecotypes of salmonid fishes. Of the genes that harbored a *cis*-QTL for either splicing or transcription, we observed a similar proportion (5/22 versus 5/24) showing overlap with LFMM hits. It is possible that splicing plays as important a role as transcription in adaptation and that the two mechanisms fulfill unique roles in adaptation.

RNA-Seq remains the most cost-effective technique for studying transcriptomes, but it is difficult quantify whole isoforms from short-read data. In this study, we used split reads to identify alternative splicing events. Advances in long-read sequencing using PacBio and Nanopore technologies are now opening new avenues for studying isoforms. For example, tens of thousands of novel isoforms have been identified even in well-studied organisms like humans, (Glinos et al. 2022; Yamaguchi et al. 2022) suggesting that these approaches will be particularly useful for studying alternative splicing in an evolutionary context.

## Acknowledgements

The authors thank Leah Hyee Ryun Lee and Yocelyn Gutierrez Guerrero for their help with bioinformatic analyses, and Yocelyn Gutierrez Guerrero and Mallory A. Ballinger for their suggestions and comments. We thank Katya L. Mack and Megan Phifer-Rixey for providing data on adaptive gene expression and genome-wide signatures of selection, respectively. This work was possible thanks to an allocation of Advanced Cyberinfrastructure Coordination Ecosystem: Services and Support (ACCESS) to Michael W. Nachman and Megan Phifer-Rixey. This work was supported by the National Institutes of Health (R01 GM127468 and R35 GM149304).

## Author contributions

DNM and MWN designed the project. DNM conducted the analyses and produced the figures. Both authors reviewed and interpreted the data and contributed to writing the manuscript.

## Competing interests

The authors declare no competing interests.

## Notes

### Competing Interest Statement

The authors have declared no competing interest.

