## Supplementary figures for "Alternative splicing and environmental adaptation in house mice"

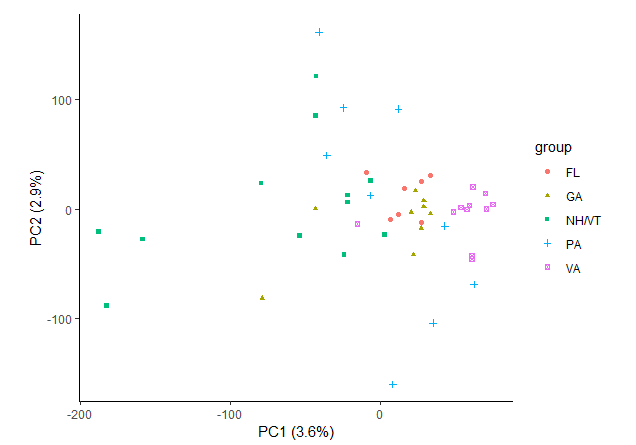

**Supplementary Fig 1. Principal component analysis of variation in splicing data.** The principal components were calculated from junction read counts across all genes.

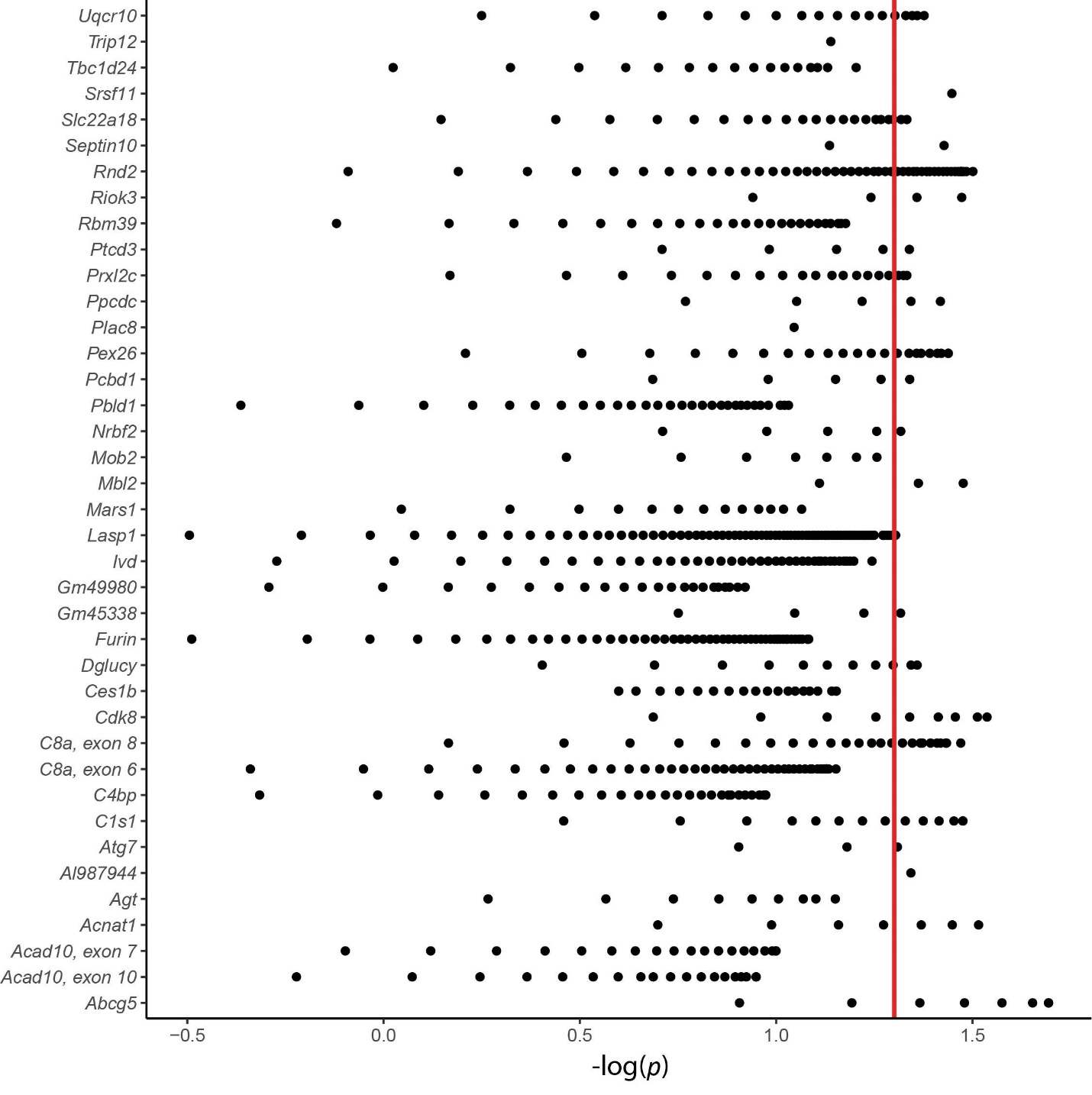

**Supplementary Fig 2. Identification of *cis*-sQTL**. For each of 44 genes found to have clinal variation in splicing, GEMMA was used to calculate the association between splice-in frequency and every SNP found within 200 kb of the respective exon’s acceptor site. Significant SNPs were classified as *cis*-sQTLs (-log_10_(*p*) > 1.30, *p* < 0.05, Benjamini-Hochberg correction). Each point represents a SNP and its significance. SNPs that fell above the threshold (red line) are *cis*-sQTL.

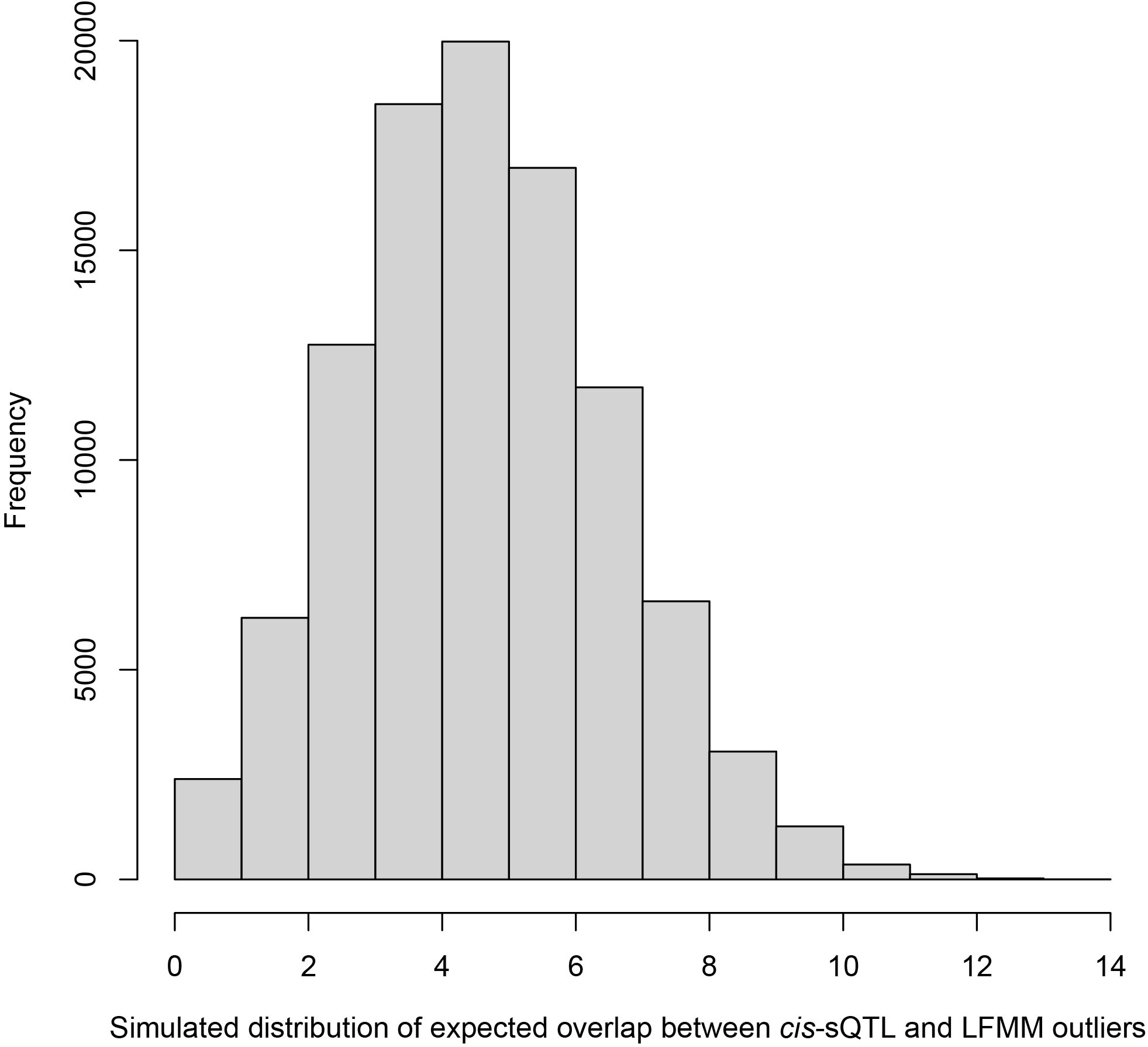

**Supplementary Fig 3. Permutation test for overlap between *cis*-sQTL and LFMM outliers.** Genes were randomly sampled from expressed genes in house mouse liver tissue 22 times (representing the 22 *cis*-sQTLs) and 5643 times (representing the number of LFMM outliers) 100,000 times.

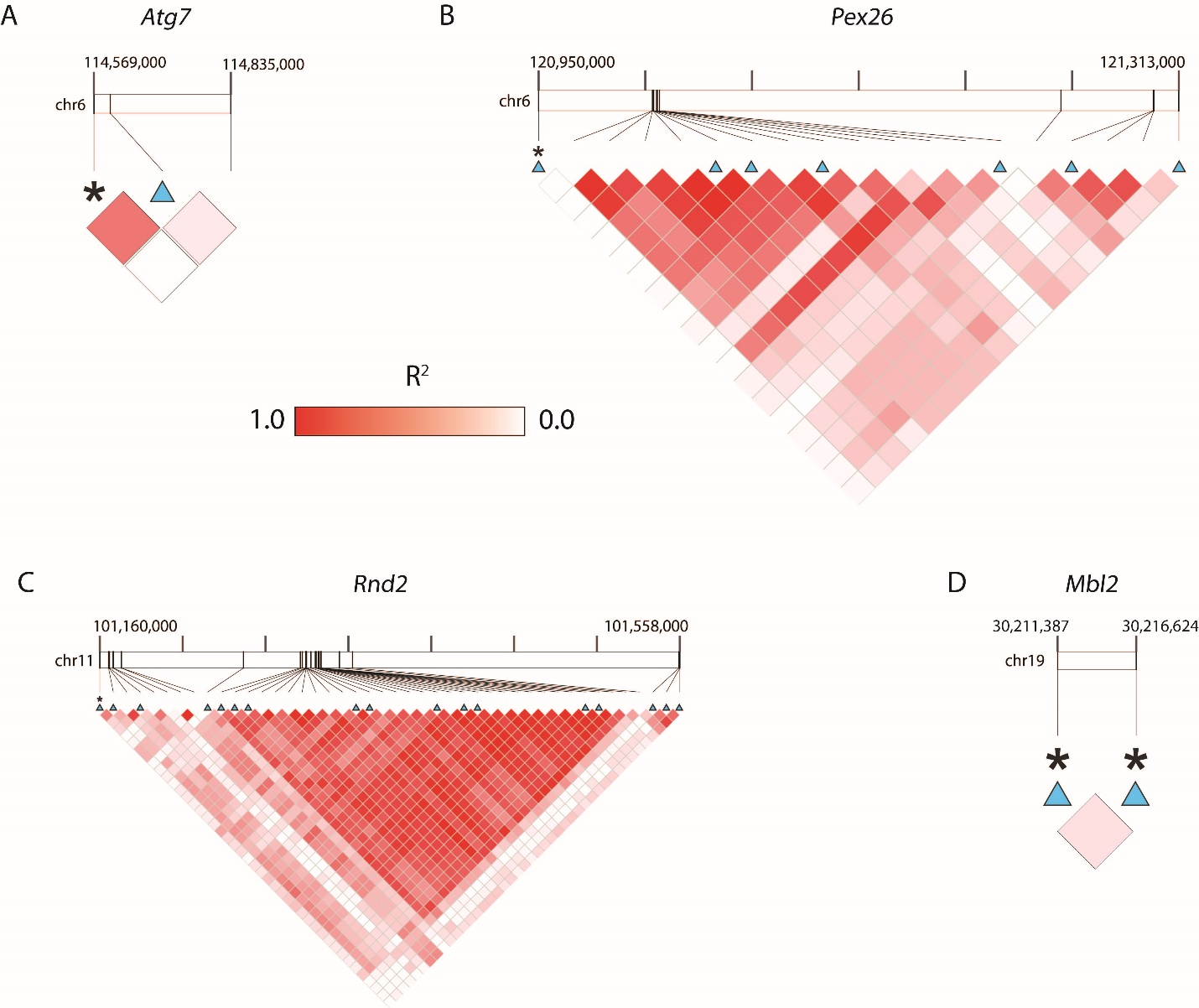

**Supplementary Fig 4. Linkage disequilibrium between *cis*-sQTL and LFMM outliers.** Linkage disequilibrium was calculated between every SNP within 200 kb of an exon acceptor site for clinally varying alternatively spliced transcripts. Asterisks indicate targets of selection (Latent Factor Mixed Model outlier SNPs) and blue triangles indicate *cis*-sQTL. *Pex26*, *Rnd2*, and *Mbl2* all had co-localized SNPs. Exome sequencing captured very few SNPs within 200kb of *Atg7* and *Mbl2*.
